# Genotype-specific expression of *uncle fester* suggests a role in allorecognition education in a basal chordate

**DOI:** 10.1101/2024.02.13.580188

**Authors:** Daryl A. Taketa, Liviu Cengher, Delany Rodriguez, Adam D. Langenbacher, Anthony W. De Tomaso

## Abstract

Histocompatibility is the ability to discriminate between self and non-self tissues, and has been described in species throughout the metazoa. Despite its universal presence, histocompatibility genes utilized by different phyla are unique-those found in sponges, cnidarians, ascidians and vertebrates are not orthologous. Thus, the origins of these sophisticated recognition systems, and any potential functional commonalities between them are not understood. A well-studied histocompatibility system exists in the botryllid ascidians, members of the chordate subphylum, Tunicata, and provides an opportunity to do so. Histocompatibility in the botryllids occurs at the tips of an extracorporeal vasculature that come into contact when two individuals grow into proximity. If compatible, the vessels will *fuse*, forming a parabiosis between the two individuals. If incompatible, the two vessels will *reject*-an inflammatory reaction that results in melanin scar formation at the point of contact, blocking anastomosis. Compatibility is determined by a single, highly polymorphic locus called the *fuhc* with the following rules: individuals that share one or both *fuhc* alleles will fuse, while those who share neither will reject. The *fuhc* locus encodes multiple proteins with roles in allorecognition, including one called *uncle fester,* which is necessary and sufficient to initiate the rejection response. Here we report the existence of genotype-specific expression levels of *uncle fester*, differing by up to 8-fold at the mRNA-level, and that these expression levels are constant and maintained for the lifetime of an individual. We also found that these differences had functional consequences: the expression level of *uncle fester* correlated with the speed and severity of the rejection response. These findings support previous conclusions that *uncle fester* levels modulate the rejection response, and may be responsible for controlling the variation observed in the timing and intensity of the reaction. The maintenance of genotype specific expression of uncle fester is also evidence of an education process reminiscent of that which occurs in mammalian Natural Killer (NK) cells. In turn, this suggests that while histocompatibility receptors and ligands evolve via convergent evolution, they may utilize conserved intracellular machinery to interpret binding events at the cell surface.

## Introduction

Histocompatibility, also known as allorecognition, is the ability of an individual to discriminate its own tissues from those of another individual and is found throughout the metazoa. This recognition event is used in a variety of processes, including the prevention of inbreeding in hermaphrodites, mediating social interactions and kin selection, and is the basis of the vertebrate adaptive immune system (Burnet 1971) . Although histocompatibility is conserved, the candidate genes identified in different species are not, suggesting that this complex recognition process has evolved independently in groups as diverse as marine sponges, cnidarians and ascidians (Grice and Degnan 2015). However, in contrast to innate immunity, where non-self epitopes are recognized by innate, germline encoded immune receptors-e.g., pathogen associated molecular patterns (PAMPS) by germline encoded pattern recognition receptors (PRRs), allorecognition is dependent on the ability to discriminate between protein alleles (allotypes) of highly polymorphic ligand(s), which requires a recognition event that is orders of magnitude more specific. The only system where the molecular mechanisms of histocompatibility are known is the Major Histocompatibility Complex (MHC) based adaptive immune system in the vertebrates, and in this case, histocompatibility requires a number of unique molecular and cellular processes, including the ability to recombine and mutate genomic DNA (somatic recombination), as well as the presence of a unique organ (the thymus), which mediates a process called thymic selection that establishes specificity of the immune reaction for each individual (Ashby and Hogquist 2024). However, neither the MHC, nor the proteins required for somatic recombination are found in non-vertebrates. In addition, to date there is no evidence that these recombination or education processes are required for histocompatibility in invertebrate species (De Tomaso 2009).

In addition to the adaptive immune system, mammals also have an innate allorecognition response mediated by Natural Killer (NK) cells, which use germline encoded receptors to detect MHC Class I molecules. In humans, these receptors are called Killer Ig-like receptors (KIRs). There are 15 KIR genes, and they are encoded in linked, diverse haplotypes that are both polymorphic and show gene content variation (Pollock et al. 2022). A subset of KIRs can discriminate between one of four polymorphic epitopes found on MHC Class I proteins, called A3/11; Bw4, C1 and C2. However, as each of these epitopes are found on multiple MHC Class I allotypes, NK-based allorecognition can only discriminate between groups of alleles, not between individual allotypes, and thus individuals. In addition, because these epitopes are not found on all MHC Class I allotypes, NK cells can also play a role in allorecognition in mammals following transplantation of MHC mismatched grafts (Kärre 2008).

The level of discriminatory ability by NK cells is much lower than both invertebrate allorecognition and the vertebrate adaptive immune system, both of which can discriminate between even closely related individuals with similar allotypes. Thus it is interesting that invertebrate allorecognition uses innate, germline encoded receptors to carry out a recognition event more similar to that of the adaptive immune system, but none of the molecules used by species in different phyla are conserved, but have evolved independently (McKitrick and Tomaso 2010). How can such complex recognition systems evolve so rapidly? And could the observed rapid evolution of the polymorphic ligands and receptors depend on the presence of other conserved processes?

The basal chordate, *Botryllus schlosseri* is an excellent model to investigate these questions. During the allorecognition reaction, finger-like projections of the extracorporeal vasculature, called ampullae, come into contact when two individuals grow into proximity. This interaction will lead to one of two outcomes. Either the two ampullae will fuse, and the two individuals will form a vascular and hematopoietic chimera, or they will reject, an inflammatory response that will result in a melanin scar, called a point-of-rejection (POR) between the two opposing ampullae, destroying the site of interaction and blocking vascular fusion (Ballarin 2012; Taketa and De Tomaso 2015). Fusion is dependent upon sharing at least one allele of a single, highly polymorphic locus called the *fuhc* (for *<u>fu</u>sion/<u>h</u>isto<u>c</u>ompatibility*; Scofield et al. 1982b; Weissman et al. 1990). If no alleles are shared, an inflammatory response will occur. The simple Mendelian genetics and ability to directly observe and manipulate the interacting tissues make *B. schlosseri* an attractive model to study the underlying molecular mechanisms of allorecognition. Furthermore, the rules of allorecognition, specifically that the sharing of a single allele signals compatibility, are analogous to the missing-self recognition found in vertebrate Natural Killer (NK) cells (described below). When ampullae from two individuals touch, they are each looking for a self *fuhc* allele on the other ampullae, and if this is not detected, a rejection ensues. Most populations of Botryllus have over 100 alleles of the *fuhc*, thus histocompatibility in Botryllus is due to an ampullae being able to discriminate a self-allele from tens to hundreds of similar non-self alleles, but how this specificity is established and maintained is not understood.

Previous genetic studies have identified six genes in the *fuhc* locus with characteristics suggesting a role in allorecognition (reviewed in Taketa and De Tomaso 2015), called: *fuhc^sec^* and *fuhc^tm^*(De Tomaso et al. 2005; Nydam et al. 2013); *fester* (Nyholm et al. 2006); *uncle fester* (McKitrick et al. 2011); *bhf* (Voskoboynik et al. 2014), and *HSP40-L* (Nydam et al. 2013). Four of these genes (*bhf*, *fuhc^sec^ fuhc^tm^* and *HSP40-L*) are polymorphic and encoded in a 109 Kb linkage group. This corresponds to a genetic distance of ca. 0.1 cM (De Tomaso et al 2005), and these polymorphisms completely correlate with fusion and rejection outcomes, suggesting that one or more of these proteins are the self-ligand, although little is known about the role of each in the allorecognition reaction (Taketa and De Tomaso 2015). *Fester* and *uncle fester* are encoded ca. 150Kb upstream of *bhf*, and similar to KIRs, are encoded in diverse haplotypes that include multiple *fester* loci with gene content variation (Nyholm et al. 2006). Based on genetics, loss-of-function assays, and *in vivo* interfering antibody experiments, *fester* (Nyholm et al. 2006) and *uncle fester* (McKitrick et al. 2011), are receptors that control the fusion and rejection responses, respectively.

Both the genomic organization and polymorphism, as well as the functional roles of the Fester proteins are similar to mammalian Natural Killer cell receptors (McKitrick et al. 2011; Nyholm et al. 2006). One role of mammalian NK cells is to monitor surface expression of MHC Class I proteins on target cells. If the proteins are expressed, the cell is left alone. However, if any MHC Class I proteins are not expressed, which can occur during infection or transformation, the cell is killed. In other words, the lack of recognition of a self-ligand is one signal that a target cell is unhealthy- and this is the basis of ‘missing-self’ recognition (Karre 2008).

Despite the differences in discriminatory ability, from a functional perspective allorecognition in *B. schlosseri* is most similar to vertebrate NK-based immunity. In mammals, missing-self recognition requires two independent signals, one which activates the cell to kill, and the other which inhibits killing, with recognition of epitopes on MHC Class I being the inhibitory step. Whether a NK cell kills a target is actually a balance of these two inputs. If the activating signal is stronger than the inhibitory signal, the target is killed-but if the inhibitory signal is stronger, the target is left alone. As discussed above, the inhibitory signal comes from KIRs that bind to epitopes found on MHC Class I proteins. In addition, there are multiple activating receptors that bind a diverse range of ligands, including proteins only expressed during infection or stress. An infection can cause upregulation of activating stress ligands and/or downregulation of inhibitory MHC Class I ligands, thus the NK cell is quantifying the level of stress via integration of activating and inhibitory inputs to decide if the target cell should be destroyed (Long et al. 2013). Given that the NK cell should only kill infected cells, and leave healthy cells alone, it is clear that the maximum activating and inhibitory signals must be maintained in a ratio that allows them to detect changes in activating and inhibitory ligands. Although not well understood, this occurs during NK cell development in a process called *NK cell education*, during which both inhibitory and activating receptors are expressed stochastically until the correct balance is established, following which the expression levels are maintained in the mature cells (Boudreau and Hsu 2018) .

Allorecognition in *B. schlosseri* works in a similar fashion. In this case the activation step causes the rejection response, while the recognition of a self-*fuhc* allele is the inhibitory signal, which overrides rejection and induces fusion. And like NK cells, the outcome is a balance of activating and inhibitory inputs. We have found that *fester* is equivalent to an NK inhibitory receptor, while *uncle fester* is analogous to a NK activating receptor. siRNA-mediated knockdown of *uncle fester* completely blocked rejection responses when incompatible knockdown colonies were paired, but had no affect on fusion of compatible knockdown pairings. In addition, *in vivo* stimulation of a single ampullae with an anti-Uncle Fester mAb induced a rejection response (McKitrick et al. 2011). Importantly, when the same antibody was placed in contact with two compatible ampullae as they began to fuse, the presence of the antibody would override the fusion response, and result in a rejection. Conversely, when an antibody to Fester is added to incompatible ampullae as they begin to reject, the presence of the antibody would override the rejection response, resulting in fusion of some of the ampullae (Nyholm et al. 2006). In summary, these results demonstrated that fusion and rejection are not mutually exclusive, but due to a balance of two signals-one controlling rejection, and one controlling fusion, equivalent to mammalian NK cell responses (Long et al. 2013).

Given these functional similarities, we have hypothesized that education processes similar to mammalian NK cells may also occur in *B. schlosseri,* which would predict differences in the activation signal. Thus, we assessed *uncle fester* expression in wild type genotypes, focusing on a group of individuals with known fusion/rejection outcomes to see if uncle fester expression correlated with histocompatibility outcomes.

## Methods

We maintained *Botryllus schlosseri* colonies as previously described (Braden et al. 2014) and reared only a single genotype in each tank to prevent genetic contamination. Samples from either whole colonies or surgically removed ampullae were homogenized to a frozen powder with a mortar and pestle on dry ice with liquid nitrogen and then stored at -80°C. Total RNA was extracted from frozen powder with the NucleoSpin® RNA kit (Macherey-Nagel, 740955). RNA samples were concentrated with NucleoSpin® RNA Clean-up XS (Macherey-Nagel, 740903) or GeneJet RNA Cleanup and Concentration Micro Kit (Life Technologies, K0841) as needed. mRNA was purified from 4 µg of total RNA with a magnetic mRNA isolation kit (New England Biolabs, S1550S). cDNA was synthesized from mRNA samples with M-MLV (New England Biolabs, M0253S) primed with random hexamers (Life Technologies, 48190-011).

For mRNA-Seq, at least 500 ng of total RNA from ampullae tissue was prepared from a given genotype and was sequenced on an Illumina Hi-Seq 2000 by the USC Epigenome Center. Reads were trimmed and mapped to a public *Botryllus* EST assembly (http://octopus.obs-vlfr.fr/public/botryllus/blast_botryllus.php, database: “Bot_asmb assembly 04.05.2011, A. Gracey”) as described in Rodriguez *et al*. (2014) and differential gene expression was analyzed using the DESeq package (version 1.10.1) in R (Anders and Huber, 2010). Four genotypes analyzed by Rodriguez *et al*. (2014) were included in the DESeq analysis (GEO accession# GSE62112) to compare ampullae tissue (n=15) versus whole colony (n=28).

Quantitative Polymerase Chain Reaction (qPCR) was performed on a LightCycler480 (Roche) using SYBR Green I master mix (Roche). Equal amounts of cDNA were used to assess relative expression of *uncle fester* compared to *ef1*α using the following exon spanning primers: *uncle fester* (exons 2 to 3) forward – 5’GGATGGCTGCACCAACTACT 3’, reverse – 5’ TCACCAACGGTCGTAGAACA 3’; *ef1*α forward – 5’ TGGATTCCTCGGTGATTCTC 3’, reverse – 5’ CGAATTTTTCGATCGCTCTC 3’. The cycling conditions were as follows: 95°C – 10 sec, 56°C – 10 sec, 72°C – 20 sec. Analysis of relative expression was calculated using the 2^-ΔΔCT^ method as previously described (Livak and Schmittgen 2001). Absolute qPCR was performed utilizing a 5-fold serial dilution across 11 wells of a mammalian expression plasmid containing exons 1-5 of *uncle fester* as a standard curve to extrapolate fM amounts of *uncle fester* transcripts.

Genomic DNA (gDNA) was extracted from frozen powder using the NucleoSpin® Tissue kit (Machery-Nagel, 740952). We utilized FPROM (Solovyev et al. 2010) and NNPP (Reese 2001) to identify the putative TATA box and transcriptional start site (TSS). Additionally, a tandem repeat region (TRR; Benson 1999) was predicted ∼1800 bp upstream of the predicted TSS. The region after the TRR to the middle of exon 1 was amplified from 50 ng of gDNA using Phusion® High-Fidelity polymerase (New England Biolabs, M0530) according to the manufacturer’s instructions with the following primers: forward – 5’ AATAATCGTAAGCATGTGTTTTCTGATGTGC 3’; reverse– 5’ GAGCTTGTGTACTATTGTACGGTAAGC 3’. Amplicons were cloned into pGEM-T-Easy (Promega, A1360) after adding an A-overhang with Taq DNA polymerase (New England Biolabs, M0273). Positive clones were selected and sequenced at the UC Berkeley Sequencing Facility.

Rejection assays were conducted by transferring incompatible naïve colonies to a microscope slide with ∼2-3 mm between them. The slide was then placed in a tank with seawater until rejection occurred. Asymmetric rejection was scored by examining the location of the PORs with respect to both the ampullae and fused tunic boundary under a dissecting microscope. In most cases, a POR formed directly adjacent to reacting ampulla. PORs that were equidistant between the two ampullae or covered a large area were scored as ‘equal’ or ‘not determined’.

## Results

We had previously found that the extracorporeal vasculature of Botryllus is highly regenerative, and following surgical ablation will regrow to its original size in 48-72 hours (Nyholm et al. 2006). We were interested in the stability of expression of different allorecognition proteins, so we took seven genotypes, surgically removed and collected the vascular tissue, allowed the vessels to regenerate, and repeated the tissue isolation 2-5 times for each genotype. mRNA was isolated from each sample and either used for mRNA-seq or qPCR. Differential gene expression analysis comparing ampullae mRNA to our previously published whole colony mRNA-Seq samples (Rodriguez et al. 2014) identified consistent genotype-specific differences in the expression level of *uncle fester* (Fig. 1). The differences were drastic, ranging from 3-to 8-fold between genotypes. We next compared global gene expression between ‘high’ and ‘low’ *uncle fester* expressing genotypes to identify genes that were expressed in a pattern similar to *uncle fester* (data not shown). However, no genes were identified, and in addition none of the other candidate allorecognition genes were found to correlate with *uncle fester* expression.

**Figure 1.**
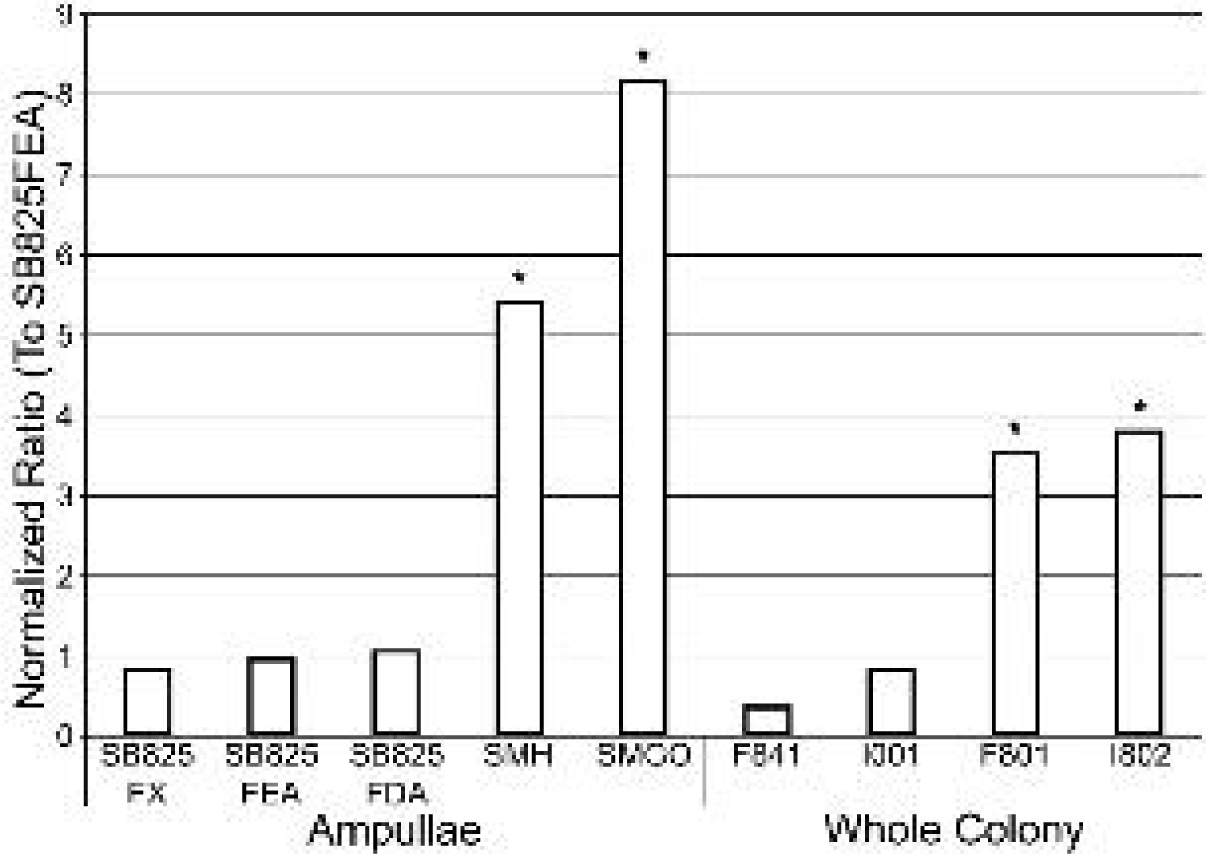
DE-Seq analysis of *uncle* fester gene expression levels in nine B. schlosseri genotype. Each DE-Seq analysis was compared against SB825FEA with the normalized ratio plotted. Asterisks represent a statistical difference (padj <0.05)

To validate our mRNA-Seq data, we quantified *uncle fester* transcripts by relative qPCR (Fig. 2b). Despite showing differences in the fold-changes between genotypes, the trends in *uncle fester* expression demonstrated by qPCR and mRNA-Seq were largely the same. In both sample populations, genotypes were identified displaying differential levels of *uncle fester* expression that were maintained in newly regenerated ampullae. Amongst the genotypes used across both analyses, the trend of a higher (e.g. SMGO) and lower (e.g. SB825) *uncle fester* genotypes remained consistent. An absolute quantification of *uncle fester* transcripts utilizing a standard curve and with equal amounts of total RNA was also performed (Supplemental Fig. 1) and also supported the differential expression of *uncle fester*. In summary, while different methodologies gave different ratios, in all cases the changes in expression were significant, and more importantly, highly repeatable, even following multiple ablation/regeneration cycles. This demonstrates that there is genotype specific mRNA expression levels of *uncle fester*.

**Figure 2.**
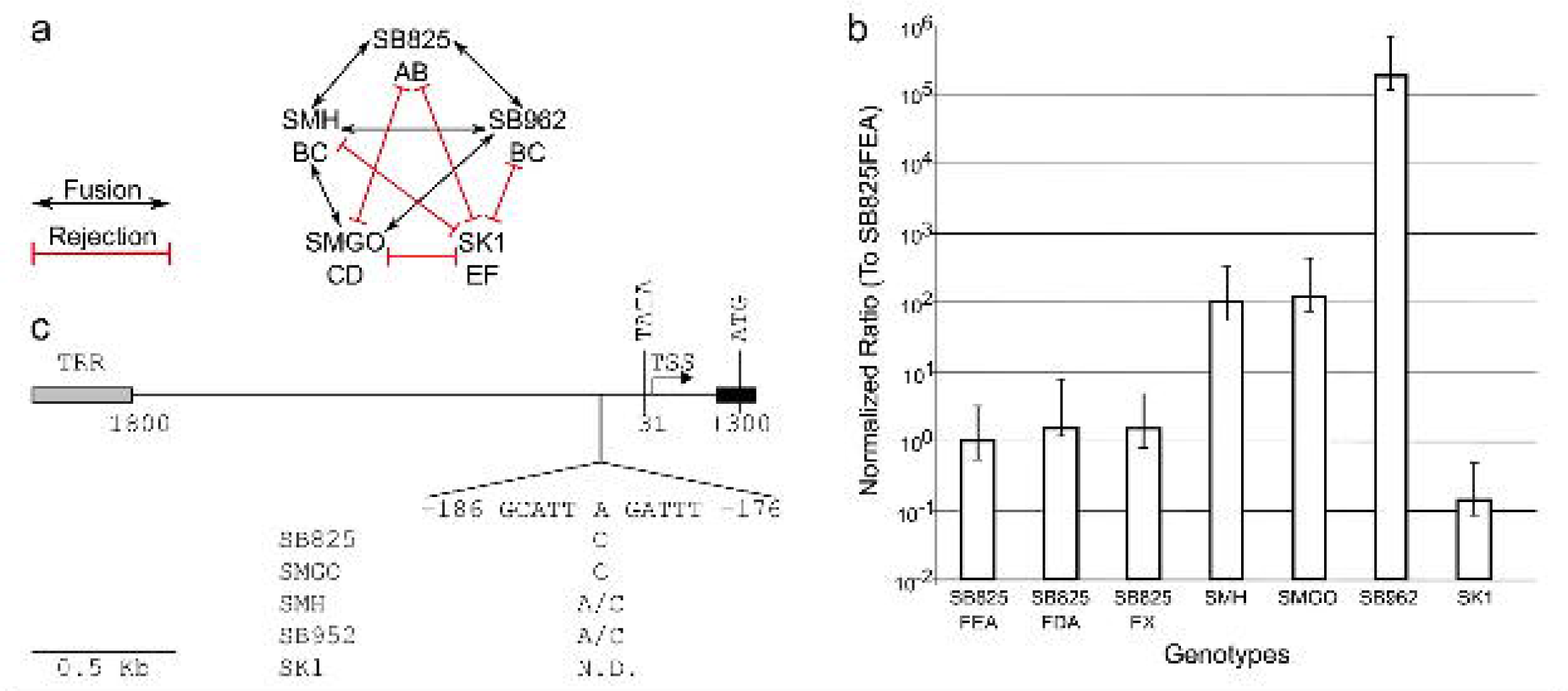
(a) Graphical representation of histocompatibility outcome amongst different lines used in this study *fuhc* alleles are denoted below the genetic line.(b) Relative qPCR analysis across several genotypes. Each sample was normalized ratio against SB825FEA was plotted with standard deviation as the error bars. Plot represents a minimum or 3 qPCR runs. (c) Graphical representation of the predicted promoter region of *uncle* fester with identified polymorphisms from each gentype listed below. Annotated posttions are relative to the TSS(+1). TRR = Tandem Repeat Region; TATA = TATA box; TSS = Transcriptional Start Site; N.D. = Not Determined

We next assessed if there were functional consequences from differences in *uncle fester* expression. Given the previously published role for *uncle fester* (McKitrick et al. 2011), we postulated that interaction between genotypes expressing different levels of *uncle fester* might produce a rejection phenotype similar to the ‘one-way’ rejection observed during knock-down experiments of *uncle fester*. This is manifested as a preferential rejection, whereby the points of rejection (POR) form near ampullae of one or the other colony (Fig. 3, white arrows). This is in contrast to equal rejection, where the POR form at the site of contact of the ampullae from both colonies (Fig. 3, black arrows). We used five wild-type genotypes and tested all combinations for fusion and rejection (Figure 2A), as well as characterizing the rejection for each pair (Table 1). In general, the genotypes expressing higher levels of *uncle fester* tended to exhibit more PORs localized near their ampullae (preferential rejection) than their partners. For example, in the pairing between SM825 and SMGO, *uncle fester* expression in SM825 was significantly lower than that of SMGO (Fig. 2B), and this correlated to the rejection response, as 84.06% of the PORs observed were preferentially located near SMGO ampullae. Interestingly, when SB962 was paired with SK1, the PORs that formed appeared to be more balanced but still had a slight preference towards SB962 (43.18% of the PORs compared to 31.82% in SK1). This bias in POR formation was less dramatic than expected given the higher expression level of uncle fester in SB962 compared to SK1. However, similar results were observed when SK1 was paired with SMH (52.17% of PORs for SMH compared to 47.83% in SK1). The SMH genotype possessed the same *fuhc* alleles as SB962 based on histocompatibility assay (Fig. 2a) and displayed a significantly much higher expression of *uncle fester* transcripts compared to SK1 (Fig. 2b). When SK1 was paired with SMGO, a genotype with similar *uncle fester* expression as SMH but differed by one *fuhc* allele, a strong bias towards SMGO was observed. One potential explanation is that differences in the *fuhc* locus, and ultimately differences in the allodeterminant protein, have an effect on the rejection phenotype (discussed below).

**Figure 3.**
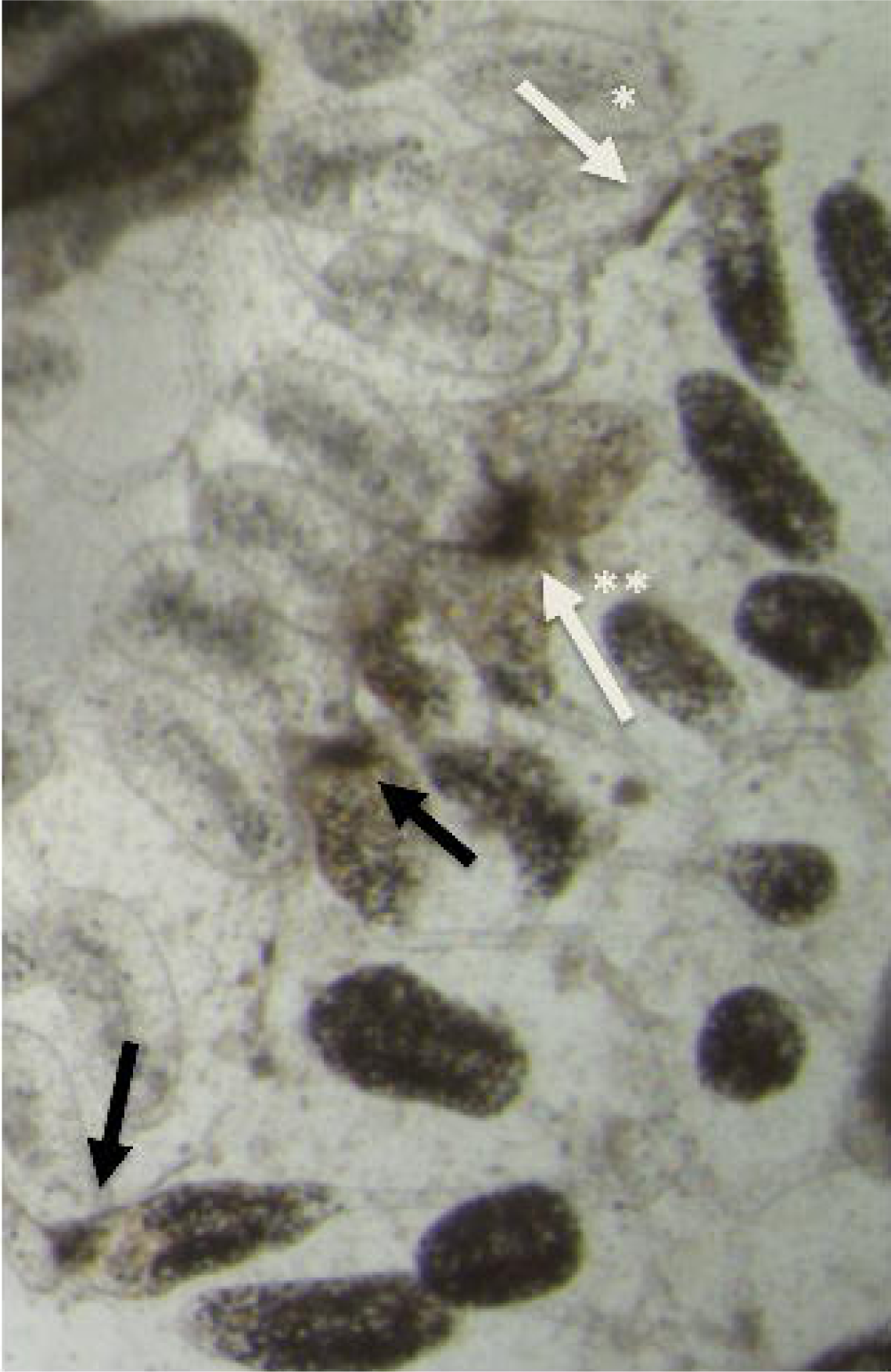
Examples of equal and preferential rejection. Interface of two incompatible individuals during a rejection response. Ampullae from top colony are clear, ampullae from bottom colony are dark. Equivalent points of rejection (POR) are highlighted with black arrows, where the two ampullae are in contact and POR forming between them. Preferential POR are highlighted with white arrows, where POR are forming from ampullae of only one of the colonies, here from either the top (*), or bottom colony(**).

**Table 1.** Preferential rejection between genotype pairs based on the numbers of Points of Rejections (PORs) and where they are formed relative to both genotypes. Examples of each are shown in Figure 3.

| Genotype 1 | Genotype 2 | % of PORs with preferential rejection in Genotype 1 | % of PORs with preferential rejection in Genotype 2 | % of PORs with equal rejection | # of PORs examined |
| --- | --- | --- | --- | --- | --- |
| SK1 | SMGO | 6.06 | 90.91 | 3.03 | 33 |
| SB825 | SMGO | 15.94 | 84.06 | 0.00 | 69 |
| SK1 | SB825 | 21.28 | 61.70 | 17.02 | 47 |
| SK1 | SMH | 47.83 | 52.17 | 0.00 | 23 |
| SK1 | SB962 | 31.82 | 43.18 | 25.00 | 34 |

Lastly, we investigated if *uncle fester* expression was regulated by any polymorphisms in the promoter region. Within the ∼1900 bp region sequenced, only a single nucleotide polymorphism (SNP), an A or C, was identified ∼181 bp upstream of the predicted TSS (Fig. 2c). However, this SNP did not genetically correlate to the expression levels of *uncle fester* or change any transcription factor binding site based on TFBIND prediction (Tsunoda and Takagi 1999). Instead, the genotype-specific expression of *uncle fester* could be attributed to the differential expression of transcription factors (Supplemental Table 1), polymorphisms in regulatory elements further upstream, or polymorphisms in the tandem repeat region.

In summary, we report that there are genotype specific expression levels of *uncle fester*, and that these expression differences are stable and maintained in newly regenerated ampullae. Amongst incompatible pairs with differential *uncle fester* expression, genotypes with the higher expression usually exhibited more PORs than their lower expressing partners. These findings parallel our previous *uncle fester* knockdown experiments, and strongly suggest that the expression level of *uncle fester* is a key modulator of the rejection response.

## Discussion

We have previously found that allorecognition in *B. schlosseri* consists of two mutually exclusive steps, each mediated by a different fester family protein. siRNA-mediated knockdown revealed that *uncle fester* is necessary and sufficient to initiate the rejection response, but had no effect on fusion. Conversely, *fester* expression is necessary for fusion (McKitrick et al. 2011; Nyholm et al. 2006). Using monoclonal antibodies, we also found that both rejection and fusion responses can be manipulated *in vivo*: stimulation of Uncle Fester following contact of ampullae of two compatible colonies caused them to reject, while stimulation of Fester following contact of ampullae of two incompatible colonies caused them to fuse. This demonstrates that each of these inputs are tunable, and that outcome (fusion or rejection) is due to the integration of these two signals (McKitrick et al. 2011; Nyholm et al. 2006).

These results are functionally equivalent to studies in mammalian Natural Killer cells, where it has been shown that outcome of an interaction with a target cell is also due to integration of activating and inhibitory inputs (Long et al. 2013), due to the presence and/or absence of ligands on the target cell surface. If the strength of activation is higher than inhibition, the target is killed, but if inhibitory strength dominates, the target cell is tolerized. Since the role of the NK cell is to use this balance to discriminate between healthy and unhealthy cells, and failure to do so would result in either a failed immune response, or autoimmunity, the range of activating and inhibitory signaling must be carefully calibrated. Calibration is accomplished via a process called NK cell education that occurs during development (Boudreau and Hsu 2018). While the education process is not well understood, it involves stochastic expression of both activating and inhibitory receptors from the receptor haplotypes until the correct balance is achieved, resulting in populations of educated cells that each have a random group of activating and inhibitory receptors. The balance between activating and inhibitory responses can be detected via analysis of these educated cells. For example, there is a positive correlation between the number of inhibitory receptors a NK cell expresses, and the maximum activation that can be stimulated (Joncker et al. 2009). In summary, NK cell education ensures that the activating and inhibitory inputs are coordinated, and this process is vital for NK immunity. To detect and integrate differences in activating and inhibitory ligand expression on target cells, the maximum activation a cell can detect cannot be higher than what can be inhibited, or those cells could be autoreactive. Conversely, if the maximum inhibition is higher than the potential activation, those cells could be immune deficient. While the cellular and molecular mechanisms underlying education are unknown, they must depend on the ability of cells to monitor and quantitate binding events on the cells surface, and compare them to a preexisting threshold value. This ability has been termed *quality control* (Boehm 2006).

As described above, in *B. schlosseri* fusion and rejection are also not mutually exclusive events, rather based on the integration of two independent signals, suggesting that same type of education process must occur. In addition, there is data that suggests this is also a stochastic process. Previous studies have demonstrated phenotypic variability in the rejection reaction-the severity of the rejection phenotype in Botryllus schlosseri is variable (Scofield and Nagashima 1983; Oren et al. 2008), and can be classified into one of four categories, depending on the time of POR formation as well as the number of PORs observed. Interestingly, forward genetic analysis of this variability mapped rejection intensity to the *fuhc* locus itself (Scofield and Nagashima, 1983; discussed below). In addition, a previous study found that in some incompatible pairings a ‘one-way’ phenotype could be observed, where ampullae from one individual had multiple POR, while ampullae from the other individual did not, and termed the responsive colony the ‘rejector’ and the unresponsive colony the ‘rejected’ (Oren et al., 2010). These observations raise the question of whether variations in rejection phenotypes (severity and directionality) are dependent upon differences in the sequence or expression level of the allodeterminant, as proposed by Scofield and Nagashima (1983), and/or the expression levels of *uncle fester*. In an earlier study, we tested if the rejection response was due to homotypic binding of uncle fester, we paired an *uncle fester* knockdown to an unmanipulated colony. We observed the ‘one-way’ rejection phenotype, whereby PORs were observed proximal to ampullae from the unmanipulated colony, but not from the *uncle fester* knockdown colony. In addition, partial knockdown of *uncle fester* also correlated with rejection severity. Together this suggests that changes in *uncle fester* expression can correlate with the rejection phenotype, and in addition that rejection is not initiated via homotypic binding of Uncle Fester (McKitrick et al., 2011).

Here we show that there are genotype specific expression levels of *uncle fester,* and that these levels are maintained over time, and following multiple ampullae ablation/regeneration cycles. Expression levels differ by orders of magnitude, and correlate with rejection severity *in vivo*. Interestingly, expression level differences did not correlate with any other gene expression patterns (Supplemental Table 1). This suggests that there is a plasticity to *uncle fester* expression during embryonic development that is then locked down in the adult body plan, and that this variability is due to the stochastic nature of an education process. We have also found evidence of education for the inhibitory gene, *fester* (Nyholm et al. 2006), and given the integration of the Fester and Uncle Fester pathways, it is clear that these two signals must also be calibrated via quality control mechanisms functionally equivalent to those in NK cells.

In summary, despite a lack of conservation of the ligands and receptors used in Botryllus allorecognition versus mammalian NK cells, both systems require the cells to have a quality control system that ensures calibration between the two inputs to ensure specificity. We have hypothesized that it is these intracellular quantitative processes mechanisms that are conserved across the metazoa (De Tomaso 2009). If these education processes had an early evolutionary origin, this could explain the rapid and convergent evolution of highly polymorphic recognition systems throughout the metazoa, as a conserved intracellular quality control process would allow cell surface ligands and receptors to evolve freely. The variability and long-term maintenance of genotype specific uncle fester expression provides a model to study these education processes.

## Acknowledgements

The authors thank Michael B. Caun and Greg Stoney for their expert care of the De Tomaso laboratory mariculture facility. This work was funded by the following grants through the National Institutes of Health: R35 GM139649 (to AWD) and F32 GM108227 (to ADL). This work was also funded by California Institute for Regenerative Medicine: T3-00009 (to DR).

## Conflict of Interest

The authors declare that they have no conflict of interest.

---

**Supplemental Table 1.** Differential expressed transcription factors that were upregulated in either high or low expressing *uncle fester* (*uf*) samples

| EST Contigs | Human Homolog | Log<sub>2</sub> Fold Change<br>(*uf* High/*uf* Low) |
| --- | --- | --- |
| Bot_c27848 | Zinc finger protein 596 | 4.295190775 |
| Bot_c5766 | General transcription factor 3C polypeptide 3 | 3.546378836 |
| Bot_rep_c40091 | Zinc finger protein 484 | 3.181889999 |
| CAP3_round1_contig_2290 | Transcription initiation factor TFIID subunit 11 | 2.391093104 |
| Bot_rep_c37237 | Poly [ADP-ribose] polymerase 14 | 2.203390801 |
| Bot_c22244 | DNA-directed RNA polymerase III subunit RPC8 | 1.842511855 |
| Bot_c8226 | Zinc finger protein 43 | 1.74109414 |
| Bot_rep_c36589 | Transcriptional repressor NF-X1 | 1.670011357 |
| CCAO1293.g1 | Transcription initiation factor TFIID subunit 5 | 1.571675768 |
| CCTT36475.g1 | Retinoblastoma-binding protein 5 | 1.488219578 |
| CCAO834.b1 | Zinc finger protein ZFAT | 1.482147469 |
| Bot_rep_c36926 | Transcriptional enhancer factor TEF-3 | 1.478247656 |
| Bot_rep_c45954 | DNA-directed RNA polymerase III subunit RPC10 | 1.474068851 |
| Bot_c7118 | Tuberin | 1.473017389 |
| CCTU1031.b1 | Zinc finger protein 845 | 1.467475577 |
| Bot_c19681 | Glucose-6-phosphate 1-dehydrogenase | 1.443424698 |
| Bot_rep_c45991 | DNA-directed RNA polymerase II subunit RPB4 | 1.239840315 |
| CAP3_round1_contig_1147 | Transcription factor CP2-like protein 1 | 1.047034394 |
| Bot_rep_c49277 | Transcription initiation factor TFIID subunit 9 | -1.085452134 |
| Bot_c5850 | Zinc finger protein 161 homolog | -1.199508143 |
| Bot_c20331 | Inhibitor of growth protein 3 | -1.199541493 |
| Bot_rep_c35663 | Small nuclear ribonucleoprotein Sm D3 | -1.23446094 |
| Bot_c28224 | Zinc finger protein 235 | -1.240521962 |
| Bot_rep_c52157 | Retinoblastoma-binding protein 5 | -1.257787501 |
| Bot_rep_c36194 | Transcription initiation factor TFIID subunit 11 | -1.263510565 |
| Bot_c21801 | DNA-(apurinic or apyrimidinic site) lyase | -1.429087253 |
| Bot_rep_c54275 | REST corepressor 2 | -1.440050289 |
| Bot_c16902 | Trimethylguanosine synthase | -1.496065734 |
| Bot_c2228 | Heterogeneous nuclear ribonucleoprotein D-like | -1.500274844 |
| Bot_rep_c41708 | DNA-directed RNA polymerase II subunit RPB2 | -1.527127595 |
| CCTT2604.g1 | Transcription termination factor 2 | -1.58225855 |
| CAP3_round1_contig_5082 | NF-kappa-B-activating protein | -1.583130971 |
| Bot_rep_c35638 | DNA-directed RNA polymerases I, II, and III subunit RPABC2 | -1.598305003 |
| Bot_rep_c39662 | Cell division cycle 5-like protein | -1.6870365 |
| Bot_rep_c49315 | Activating signal cointegrator 1 complex subunit 3 | -1.690143867 |
| Bot_rep_c54145 | C-Myc-binding protein | -1.773627621 |
| Bot_c14258 | Helicase-like transcription factor | -1.812234146 |
| Bot_c19191 | Thyroid receptor-interacting protein 11 | -1.892117186 |
| Bot_rep_c36207 | 5'-3' exoribonuclease 2 | -1.892886586 |
| Bot_c17013 | FACT complex subunit SSRP1 | -1.899964921 |
| Bot_rep_c34994 | Polyubiquitin-C | -1.983413043 |
| Bot_c10893 | General transcription factor II-I repeat domain-containing protein 2B | -1.985638124 |
| CAP3_round1_contig_609<br>5 | Poly [ADP-ribose] polymerase 14 | -2.103414904 |
| Bot_c17707 | Erythroid transcription factor | -2.125455604 |
| Bot_c8616 | Putative zinc finger protein 705G | -2.143869776 |
| Bot_rep_c50646 | Heterogeneous nuclear ribonucleoprotein K | -2.320235882 |
| Bot_c21391 | Poly [ADP-ribose] polymerase 1 | -2.348468773 |
| Bot_c21036 | Polycomb protein EED | -2.427685264 |
| Bot_rep_c40327 | Poly [ADP-ribose] polymerase 14 | -2.613947182 |
| Bot_c23001 | Calcium-responsive transactivator | -2.638354453 |
| Bot_rep_c56276 | Pre-mRNA cleavage complex 2 protein Pcf11 | -2.88587826 |
| CAP3_round1_contig_313<br>5 | Heterogeneous nuclear ribonucleoprotein U-like protein 1 | -2.928125633 |
| Bot_c14707 | Zinc finger protein 583 | -2.979951203 |
| Bot_c31578 | Fez family zinc finger protein 2 | -3.321783326 |
| Bot_c20965 | Zinc finger protein 134 | -4.261153637 |

## Notes

### Competing Interest Statement

The authors have declared no competing interest.

